# Convergent genes shape budding yeast pericentromeres

**DOI:** 10.1101/592782

**Authors:** Flora Paldi, Bonnie Alver, Daniel Robertson, Stephanie A. Schalbetter, Alastair Kerr, David A. Kelly, Matthew J. Neale, Jonathan Baxter, Adele L. Marston

## Abstract

The 3D architecture of the genome governs its maintenance, expression and transmission. The conserved ring-shaped cohesin complex organises the genome by topologically linking distant loci on either a single DNA molecule or, after DNA replication, on separate sister chromatids to provide the cohesion that resists the pulling forces of spindle microtubules during mitosis^1,2^. Cohesin is highly enriched in specialized chromosomal domains surrounding centromeres, called pericentromeres^3-7^. However, the structural organisation of pericentromeres and implications for chromosome segregation are unknown. Here we report the 3D structure of budding yeast pericentromeres and establish the relationship between genome organisation and function. We find that convergent genes mark pericentromere borders and, together with core centromeres, define their structure and function by positioning cohesin. Centromeres load cohesin and convergent genes at pericentromere borders trap it. Each side of the pericentromere is organised into a looped conformation, with border convergent genes at the base. Microtubule attachment extends a single pericentromere loop, size-limited by convergent genes at its borders. Re-orienting genes at borders into a tandem configuration repositions cohesin, enlarges the pericentromere and impairs chromosome biorientation in mitosis. Thus, the linear arrangement of transcriptional units together with targeted cohesin loading at centromeres shapes pericentromeres into a structure competent for chromosome segregation during mitosis. Our results reveal the architecture of the chromosomal region within which kinetochores are embedded and the re-structuring caused by microtubule attachment. Furthermore, we establish a direct, causal relationship between 3D genome organization of a specific chromosomal domain and cellular function.

## Main text

To map pericentromere domains, we arrested cells in metaphase either in the presence or absence of microtubules and analysed cohesin (Scc1) localization by calibrated ChIP-Seq. While cohesin peaks on chromosome arms were comparable in both conditions, signal was reduced over ∼15kb surrounding centromeres in the presence of microtubule-dependent spindle tension, as reported^4,8,9^ (Figure 1a). The Wpl1/Rad61 protein promotes cohesin turnover prior to metaphase^10^, but was dispensable for the tension-dependent reduction in pericentromeric cohesin (Figure S1), suggesting that removal occurs passively. Interestingly, however, prominent peaks flanking both sides of centromeres persisted in the presence of tension, and additional small peaks appeared further away from some centromeres (Figure 1a, asterisks). Pericentromeric cohesin enrichment is achieved by the specific targeting of cohesin loading to the centromere by a direct interaction between the Ctf19 inner kinetochore subcomplex and the Scc2/Scc4 cohesin loader^11,12^. Current models posit that cohesin accumulates at sites distinct from those at which it is loaded^13^. Indeed, abolishing kinetochore-driven cohesin loading (by deletion of *CHL4*^7^, encoding a Ctf19 complex component), diminished the prominent cohesin peaks flanking centromeres (Figure 1b), suggesting that some cohesin loaded at centromeres slides bidirectionally and collects at these regions. We henceforth denote these centromere-flanking regions that retain high levels of cohesin under tension and mark the limits of the pericentromere as “borders”. Aligning pericentromere borders from all 16 chromosomes, using the centre of the first cohesin peak that persists under tension, confirmed that while cohesin at centromeres is generally diminished under tension, cohesin at borders is not, and that Chl4 promotes cohesin association with both locations (Figure 1c).

**Figure 1.**
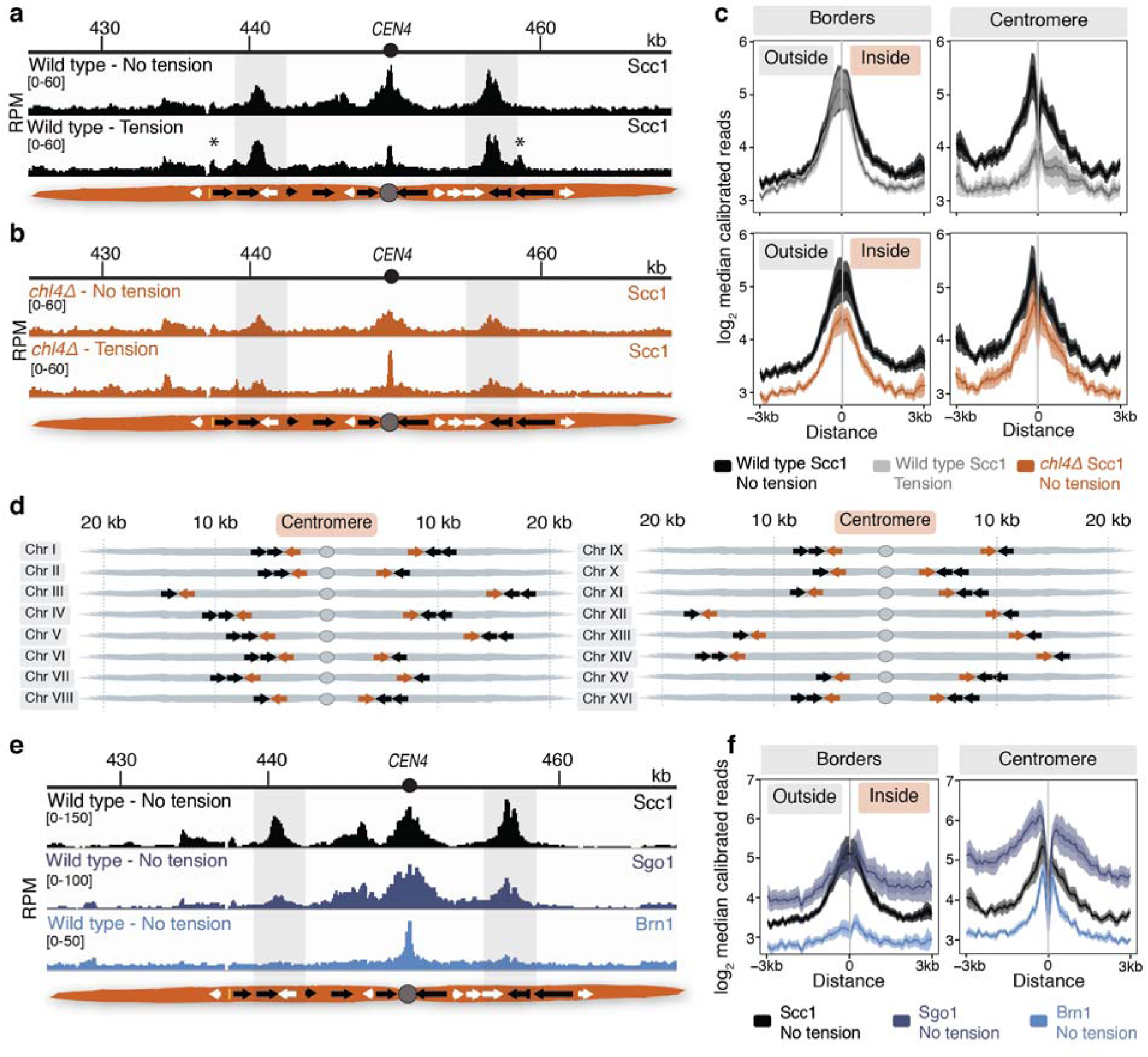
Convergent genes mark pericentromere borders. Cohesin (Scc1) enrichment in the pericentromeric region of chromosome IV in wild type (**a**) and *chl4Δ* (**b**) cells arrested in metaphase either in the presence (no tension) or absence (tension) of nocodazole and benomyl. Pericentromere border regions are shaded in grey. Black and white arrows indicate genes transcribed towards and away from the centromere, respectively. Asterisk indicates additional cohesin peak under tension. **c**, Plots show median calibrated ChIP reads (solid line) and standard error (shading) at borders and centromeres of all chromosomes for wild type and *chl4Δ*, either in the presence or absence of tension. **d**, Schematic shows the positions of convergent gene pairs flanking the centromere. Grey ovals represent the centromere, convergent gene pairs at the borders are indicated by arrows. **e**, Cohesin (Scc1), shugoshin (Sgo1) and condensin (Brn1) enrichment in metaphase-arrested cells in the presence of nocodazole in the pericentromeric region of chromosome IV is shown. **f**, Median enrichment of Scc1, Sgo1 and Brn1 around centromeres and borders in the absence of tension is shown.

Closer inspection of pericentromere borders on all chromosomes revealed the presence of convergent gene pairs, known sites of cohesin accumulation^13^, typically symmetrically arranged around the centromere (Figure 1d). Pericentromere size, as measured by distance between borders, ranges from 9.7 kb (chromosome II) to 29.8 kb (chromosome III) with a mean of ∼17 kb and does not correlate with chromosome size (Figure S2a, b). To determine whether centromere-flanking convergent gene pairs have special properties that enable them to act as cohesin-trapping borders or whether any convergent gene pair has the potential for border function, we analysed a yeast strain where the endogenous centromere (*CEN3*) on chromosome III has been removed and an ectopic centromere (*CEN6*) inserted at a chromosomal arm region^14^. As expected, absence of endogenous *CEN3* led to a loss of cohesin enrichment at the endogenous pericentromere, including at the border regions, together with the tension-sensitive accumulation of cohesin surrounding ectopic *CEN6* on the arm of chromosome III (Figure S3). Interestingly, convergent gene pairs surrounding the ectopic centromere showed increased cohesin enrichment that persisted under tension, similar to endogenous pericentromere borders (Figure S3).

The pericentromeric adaptor protein, shugoshin (Sgo1) promotes sister kinetochore biorientation and proper chromosome segregation, in part by recruiting the chromosome-organising protein condensin to pericentromeres^15,16^. Indicating that biorientation has occurred, Sgo1 dissociates from chromosomes in a tension-dependent manner^9^. Pericentromere borders show enrichment for both Sgo1 and condensin (Brn1) (Figure 1e, f), and condensin at borders, but not core centromeres, is dependent on Sgo1 (Figure S4a, b). Moreover, tension-sensitive Sgo1 resides at borders, but not core centromeres (Figure S4c, d). This implies the existence of two pools of shugoshin and suggests that pericentromere borders may elicit the signal that indicates tension-generating biorientation has been achieved.

Paradoxically, despite the high levels of cohesin, the attachment of sister kinetochores to opposite poles at metaphase of mitosis causes the separation of sister centromeres, but not chromosomal arms^17-19^. If borders define the limits of the pericentromere by trapping cohesin to resist the separation of sister chromatids at metaphase, then fluorescent *tetO/*TetR-GFP markers within the pericentromere are expected to split into two foci at metaphase, while markers outside the border are more likely to appear as a single focus (Figure 2a). We selected two pericentromeres for further analysis: chromosome I, with its clearly delineated border cohesin peaks indicating a small (13.1kb) pericentromere, and chromosome III, with less defined tension-insensitive cohesin peaks, inferring a large pericentromere (∼29.8 kb) (Figure 2b). This expected difference in pericentromere size predicts differential behaviour of GFP foci integrated at equivalent distances from the centromere. Indeed, while a GFP marker 12kb from *CEN1* was almost always observed as a single focus at metaphase, a marker 12kb from *CEN3* frequently split into two foci and, for *CEN3*, >95% cells with single foci were only observed when a marker was positioned 23kb away (Figure 2b). Measurement of inter-foci distances confirmed these findings (Figure 2c). However, this analysis also suggests stochasticity in the extent of pericentromere separation in metaphase. Although located outside the annotated pericentromere, the marker 18 kb from *CEN3* splits in ∼10% of cells (Figure 2b, c), indicating that the prominent border cohesin peak does not provide a fail-safe barrier to separation. Similarly, on chromosome I, the second peak of cohesin that persists in the presence of tension appears to play the predominant role in border function because a marker at 7kb separates in ∼58% of cells, while a marker at 8kb, within a second, distal cohesin peak, separates in only ∼30 % of cells, (Figure 2b, c). Overall, these findings suggest that, while preferred pericentromere borders exist, alternative sites of cohesin accumulation lead to cell-to-cell variability in the extent of sister chromatid separation at metaphase.

**Figure 2.**
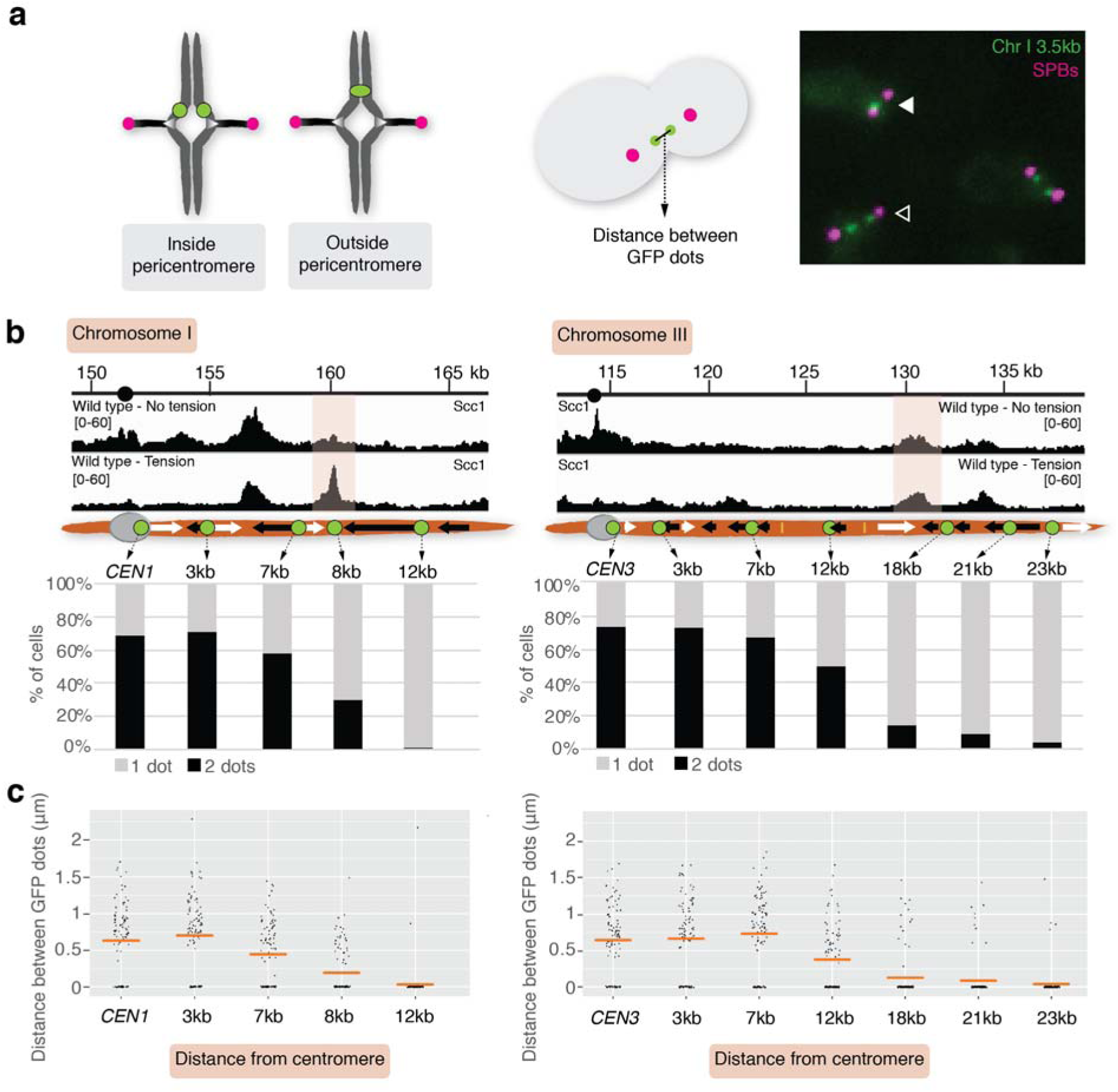
Pericentromere borders resist sister chromatid separation under tension. **a**, Assay to measure separation of loci on sister chromatids in metaphase arrested cells. Cells carry *tetO/*TetR-GFP foci integrated at various positions, Spc42-tdTomato foci to mark spindle pole bodies and are arrested in metaphase by Cdc20 depletion. Left schematic shows expected separation of GFP foci positioned inside and outside pericentromere loci. Green dots, *tetO/*TetR-GFP foci, Red dots, spindle pole bodies. Representative image and schematic of distance measured is shown to the right. White and black arrows mark cells with a single GFP focus or split foci, respectively. **b**, The number of cells with 2 GFP foci were scored for the markers at the indicated loci on chromosome I (left panels) or III (right panels) (*n* = 100). Position of GFP foci and corresponding calibrated Scc1-6HA ChIP-seq profiles are shown for comparison. **c**, The distance between GFP dots were measured (*n* = 100). Horizontal lines indicate mean.

Our data suggest that the ability to trap cohesin at border regions flanking centromeres defines the chromosomal domain that will separate under tension, which we hypothesise defines the structure of the pericentromere. A previous 3C study observed contacts between the left and right flanking regions of pericentromere III and it was suggested to be organised into an intra-molecular loop, extending between 11.5kb and 25 kb^20^. Although this predicted pericentromere size is consistent with our mapping and functional analysis (Figure 1a, Figure 2b, c), the role of borders remains unclear. To determine pericentromere structure globally and the effect of spindle tension, we performed high resolution Hi-C analysis on metaphase-arrested cells both in the presence (no tension) and absence (tension) of microtubule poisons to allow capture of unbiased genome-wide interactions. In the absence of tension at metaphase, and consistent with cis-looping in mitosis^21-23^, centromere-centered pile-up contact maps of all chromosomes showed a high frequency of cis contacts along chromosome arm regions with core centromeres acting as strong insulators (Figure 3a, first panel). The lower than expected frequency of contacts between the left and right side of the centromere (Figure 3a, second panel) argues against the presence of the previously proposed single intramolecular loop across both sides of the pericentromere^20^. Instead, close examination of individual pericentromeres or pile-ups revealed that each side of the core centromere made frequent contacts with the pericentromere on the same side, extending as far as the border 5-10kb away (Figure 3a, third and fourth panel, Figure S5). This characteristic Hi-C stripe protruding from the core-centromere is suggestive of extrusion of a chromatin loop by a centromere-anchored factor^24^. There is also evidence of longer (20-30kb) *cis* looping emanating from directly adjacent to the core centromere into either chromosome arm (Figure 3a, Figure S5). This is consistent with the notion that the usage of convergent gene pairs as boundaries is somewhat stochastic (Figure 2b, c). Interestingly, the strongest Hi-C signal occurs where there is the greatest average cohesin density at pericentromere borders (Fig 3a, third and fourth panel, Scc1 traces). In contrast, pericentromeric condensin does not appear to play an important role in pericentromere structure in the absence of tension. Hi-C maps of *sgo1Δ* which reduces pericentromeric condensin or *sgo1-3A*, which although failing to bind PP2A, recruits condensin normally^15,16^, showed pericentromeric structures that were virtually indistinguishable from wild type (Figure S6a, b).

**Figure 3.**
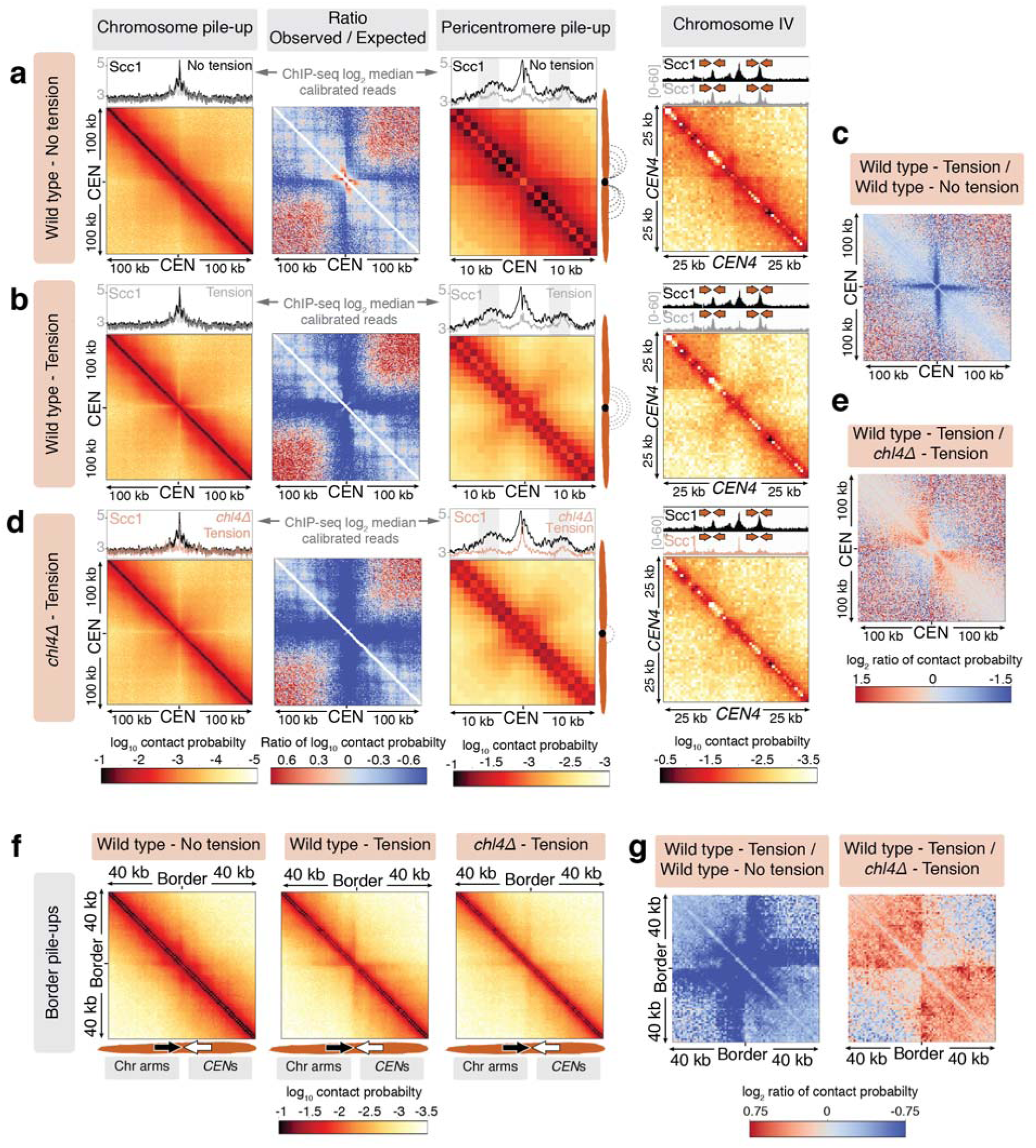
The pericentromere is a multi-looped structure in mitosis which extends to an open V shape under tension. Hi-C analysis of wild type and *chl4Δ* strains arrested in metaphase. **a, b**, Pile-ups (bin size 1kb) of *cis* contacts 100kb surrounding all 16 centromeres (left panel), ratio of expected/observed signal (second panel), pericentromere pile-up (third panel, 10kb surrounding centromeres) and contact map for pericentromere IV is shown for wild type cells in the absence (**a**) or presence (**b**) of spindle tension. Median calibrated Scc1-6HA ChIP signal around all centromeres (first and third panels) or signal for chromosome IV (fourth panel) is shown above. **c**, Log2 difference between 100kb pile-ups centered on the centromere in wild type cells in the absence and presence of tension. **d**, Maps as in (A and B) for *chl4Δ* in the presence of spindle tension. **e**, Log2 difference map comparing wild type and *chl4Δ* in the presence of spindle tension. **f**, Pile-ups (1kb bins) and **g**, ratio maps of *cis* contacts surrounding pericentromere borders (40 kb) in the indicated conditions are shown.

The presence of spindle tension changed the conformation of pericentromeres radically, while chromosome arm conformation was unchanged (Figure 3b, first and second panels). Under tension, the centromeres were no longer the point of chromosome arm insulation and instead border regions formed chromosomal arm loop boundaries ∼ 5-10kb from the core centromeres (Figure 3b, third and fourth panel). Inside the borders, the frequency of contacts within, and reaching out of, pericentromeres, was substantially reduced with a new conformation definable (Figure 3c). Contacts across individual centromeres describe an open loop or V-shaped structure with the core centromere at the apex and the borders at the tips (Figure 3b, fourth panel; Figure S5). Borders mark the boundary between the pericentromere loop and the cis-loop chromosome arm conformation.

To determine whether cohesin is required for this boundary function at borders we analysed *chl4Δ* cells, which fail to load cohesin at centromeres, leading to reduced cohesin enrichment at pericentromere borders (Figure 1c). Hi-C maps of *chl4Δ* metaphase cells in the presence of spindle tension revealed a loss of both boundary function at the borders and the strength of centromere-proximal loops (Figure 3d, e) This is consistent with the increased distance between sister centromeres at metaphase in *chl4Δ* cells^7^. These conclusions were confirmed by inspection of individual pericentromeres (Figure S5). Border regions we identified on chromosomes I (Figure 2b) and IV (Figure 1a) showed clear boundary function separating the pericentromeric domain from chromosome arms, persisting under tension but diminishing in *chl4Δ* (Fig 3a, b, d, fourth panels, Figure S6). Centering the pile-ups on the borders themselves revealed strong isolation of domains proximal and distal to the centromere, which sharpens under tension (Figure 3f, g), confirming the boundary function of borders and the dependence on *CHL4*. There is also evidence of loop extrusion^24^ at borders.

What is the property of border regions that enables the structural organization of the pericentromere? Since cohesin localization is altered by transcription^25,26^, we hypothesized that convergent transcription of border gene pairs leads to cohesin retention which results in robust inter-sister chromatid linkages that isolate domains and resist spindle forces. Indeed, RNA levels corresponding to convergent genes at borders show a narrower RNA-Seq density distribution compared to all genes, suggesting moderate expression on average (Figure S7a). Analysis of transcriptome-wide RNA pol II binding site data^27^ further revealed that active transcription at convergent gene pairs is typically higher towards, rather than away from, the centromere (Figure S7b). Outside many borders, an additional gene was oriented towards the centromere and may compensate for low centromere-directed transcription of the first gene (Figure S7b). Consistent with previous reports that transcription leads to cohesin translocation^25,26^, insertion of a *URA3* cassette between convergent genes at the left border on chromosome IV led to re-distribution of cohesin in the direction of transcription (Figure S7c).

If directional transcription at borders defines pericentromere boundaries, re-orienting convergent genes pairs into a tandem arrangement might affect pericentromere behaviour. We engineered a strain in which convergent gene pairs, together with outer centromere-facing genes at the borders of chromosome IV, are arranged into a tandem orientation, transcribing away from the pericentromere (Figure 4a). In contrast to wild type cells, the reoriented chromosome IV lost cohesin peaks at chromosome borders, while additional cohesin peaks emerged further away from the centromere, potentially forming new border regions. Furthermore, both Sgo1 and condensin (Brn1) associate with the “new” borders only on reoriented chromosome IV (Figure 4b).

**Figure 4.**
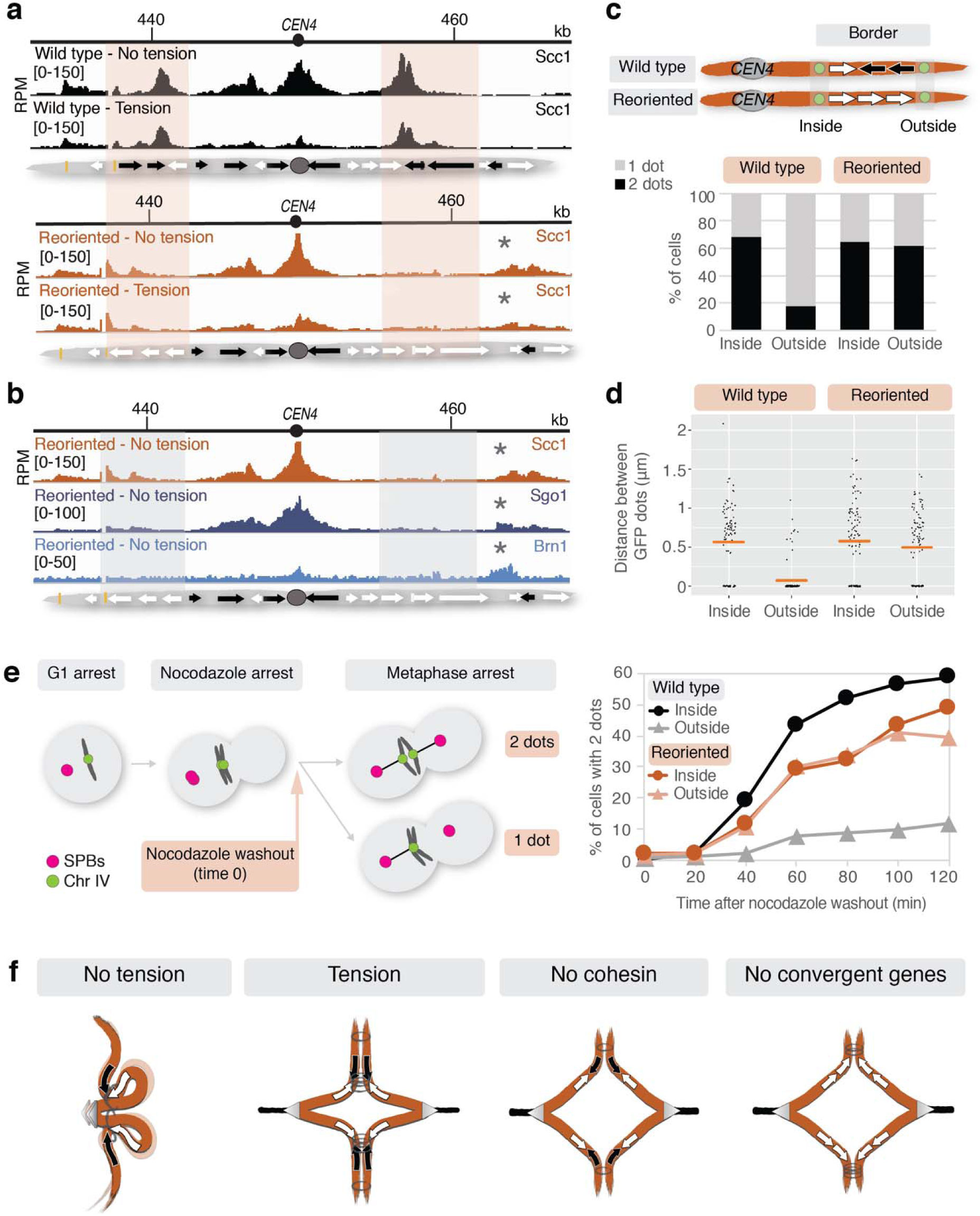
Gene orientation determines pericentromere size. The orientation of 4 genes at pericentromere borders on chromosome IV was reversed to generate a strain where both left and right borders have genes in tandem (“reoriented”). **a**, Cohesin enrichment in the pericentromeric region of chromosome IV in wild type and the reoriented strain. Shading indicates the position of the pericentromere borders in wild type cells. Asterisks indicate the position of new cohesin peaks in the reoriented strain. Schematics below show gene orientations: black and white arrows indicate genes transcribed towards and away from the centromere, respectively. **b**, Sgo1 and Brn1 localization to borders is lost following gene reorientation. Asterisks indicate new peaks in reoriented strain. **c**, Convergent genes at pericentromere borders resist sister centromere separation at metaphase. Strains with *tetO* arrays integrated at the indicated positions were arrested in metaphase and the percentage of cells with 2 GFP foci were scored (*n* = 200). **d**, Measurement of distance between GFP foci for the experiment shown in **c** (*n* = 100). **e**, Sister kinetochore biorientation following spindle re-polymerisation. Cells carrying the indicated chromosomal GFP labels, Spc42-tdTomato and *pMET-CDC20* were released from a G1 arrest and arrested in metaphase in the presence of nocodazole by depletion of Cdc20. Nocadazole was washed out while maintaining metaphase arrest by treatment with methionine and the percentage of cells with 2 GFP foci was scored at the indicated time points (*n* = 200). **f**, Model of how cohesin loaded at centromeres and convergent genes at pericentromere borders may organize the pericentromere in the presence or absence of spindle tension. For details, see text.

Since orienting the original border genes in tandem orientation causes regions more distant from the centromere to take on the role of borders, the size of the pericentromere is expected to increase. Consequently, the centromere-proximal region in which sister chromatids separate at metaphase would expand since cohesin-dependent barriers at the original border will be absent. To test this prediction, we integrated *tetO* arrays on either side of the original border in both the wild type and reoriented chromosome IV strain. As expected, GFP markers on the centromere-proximal side of the original border were separated in the majority of metaphase-arrested cells of both strains (Figure 4c, d). Remarkably, although a GFP marker outside the original border infrequently separated in wild type, in the reoriented chromosome IV strain it separated to a similar extent to a marker inside the original border (Figure 4c, d). Therefore, convergent genes set the boundaries at pericentromere borders and define the extent of sister chromatid separation at metaphase.

To determine the functional importance of pericentromere boundaries in chromosome segregation we assayed sister chromatid biorientation as the metaphase spindle re-forms after washing out microtubule-depolymerising drugs. Compared to wild type chromosome IV, reoriented chromosome IV showed a delay in, and reduced frequency of, sister kinetochore biorientation (Figure 4e). Therefore, structural organisation of the pericentromere by convergent gene pairs at borders is critical for the proper attachment of chromosomes to microtubules.

Our findings show that directed cohesin loading at a specific site coupled with cohesin stalling at distant sites together shape a chromosomal domain into a specific folded conformation (Figure 4f). Targeted cohesin loading at centromeres, and trapping between convergent genes at borders, specifically fold the budding yeast pericentromere into a multi-looped structure. We find evidence that this conformation is the product of loop extrusion on each side of the pericentromere, with borders acting to restrict loop size. This isolates each centromere from its two flanking pericentromeric regions, providing structural integrity to support the establishment of sister kinetochore biorientation. The resultant pulling forces extend pericentromeric chromatin outwards until cohesin stalling by convergent transcription at borders prevents further unzipping of the sister chromatids. In the absence of either convergent transcription (reoriented yeast strain) or efficient cohesin loading at centromeres (*chl4Δ*), borders are unable to provide the robust cohesion to resist pulling forces at metaphase and further unzipping occurs (Figure 4f).

The suggestion that cohesin makes intramolecular linkages between two sides of the pericentromere^20^ is difficult to reconcile with the strong isolation of these regions in the absence of tension (Figure 3) or the observation that cohesin is passively removed within the pericentromere when tension is applied (Figure 1a, Figure S1). Instead, we favour the idea that while some pericentromeric cohesin entraps sister chromatids to provide cohesion, other cohesin molecules make intramolecular interactions on either side of the centromere to extrude single chromatid loops. While spindle forces will pull chromatin through the sister-chromatid-entrapping cohesin until they are trapped by the transcriptional machinery at borders, intramolecular loop-extruding cohesin will be evicted from the chromosomes, consistent with the observed passive removal (Figure S8).

We have shown that a specific locus that directs cohesin loading collaborates with the linear organisation of genes to fold a chromosomal domain into a structure competent for chromosome segregation. Non-coding transcription and enrichment of cohesin are common features of centromeric regions in many organisms, suggesting general principles may underlie their structure^28^. Potentially, the linear order of transcriptional units throughout a genome has evolved in such a way to broadly influence its function by locally controlling its architecture.

## Methods

### Yeast strains and plasmids

All yeast strains were derivatives of w303 and are listed in Table S1. Plasmids generated in this study are listed in Table S2. For calibrated ChIP-Seq we used *Schizosaccharomyces pombe* strain spAM635 (h*- rad21-6HA::KanMX6*). The yeast strain carrying chromosome III with an ectopic centromere was described previously^5^. To visualize chromosomal loci, *tetO*s were integrated at defined sites on chromosome I, III and IV after cloning of the appropriate region into *pRS306(tetOx224)* (Table S2). *URA3* was inserted between convergent gene pairs by a PCR-directed approach. To reorient potential border genes on chromosome IV, the gene cassette including its promoter were cloned into a plasmid (Table S2), upstream of *KanMX*, flanked by *LoxP* sites. Plasmids were used a template for PCR, which was used for transformation, to insert the gene and its promoter in the opposite orientation, together with *LoxP-KanMX6-LoxP*. Insertion in the desired orientation was confirmed by PCR. The marker was then excised by Cre-mediated recombination.

#### Growth conditions

Cells carrying *pMET-CDC20* were arrested in metaphase in the presence and absence of tension as described by^9^. Briefly, cultures were arrested in G1 in synthetic medium lacking methionine (SC/-Met/D) with alpha factor (5 μg/ml) for 1.5 h, before re-adding alpha factor to 2.5 μg/ml and shaking for a further 1.5 h. Cells were washed with rich medium lacking glucose (YEP) and released into rich medium containing 8 μM methionine (YPDA/Met). Methionine was re-added at 4 μM every hour. To achieve a metaphase arrest in the absence of microtubules (no tension), cells were released from G1 into medium YPDA/Met containing 15 μg/ml nocodazole and 30 μg/ml benomyl. Nocadazole was re-added at 7.5 μg/ml every hour. For both the tension and no tension (nocodazole) condition, cells were harvested 2h after release from G1. For biorientation assays cells were arrested in the absence of tension as above, after 2h nocodazole was washed out by filtering with rich medium lacking glucose and cultures were released into YPDA + Met to allow spindles to reform while maintaining the metaphase arrest. Samples were taken at 20 min intervals and scored blind. To arrest cells lacking the *pMET-CDC20* construct in metaphase in the absence of spindle tension cycling cells (OD600=0.2) were treated with 15 μg/ml nocodazole and 30 μg/ml benomyl; after 1h, 7.5 μg/ml nocodazole was added and cells were harvested after a total of 2h.

#### Chromatin immunoprecipitation, ChIP-Seq and data analysis

ChIP-qPCR and ChIP-Seq was carried out as described previously^15^ except, for ChIP-Seq, purified chromatin was recovered using a PCR purification kit (Promega). Sequencing libraries were generated using standard methods and samples were sequenced on a MiniSeq instrument (Illumina) with the exception of data shown in Figures S3 and S4 where libraries were prepared and sequenced by the EMBL Genomics Core Facility. ChIP-Seq data used to generate Figure S4a was published previously^15^. Scripts, data files, and workflows used to analyse the data and prepare the ChIP-Seq figures can be found on the github repository at https://github.com/AlastairKerr. For the strains where the centromere was repositioned or where gene orientation at pericentromere borders is reversed we assembled the corresponding genome reference sequence *in silico* and used the appropriate reference to map sequencing reads for each strain. Plots showing averages of all centromeres were generated using Seqplots^29^. Read counts were normalized to reads per million mapped (RPM) and the ratio of ChIP reads to input was calculated. The mean or the median value was determined for all 16 chromosomes per 50bp window and its log2 value is graphed. Mean values are shown for the +/-3kb plots; the +/−25kb plots use median values. To allow quantitative comparison between different conditions all ChIP-Seq, with the exception of the data shown in Figure S3 and S4, was calibrated with an internal reference by modifying the procedure described by^30^. Rather than *Candida glabrata, S. pombe* carrying Rad21-6HA was used as the calibration genome (strain spAM635). Briefly, for each IP, 100 ml of *S. pombe* cells were grown in YES to OD595=0.25–0.3 and fixed by addition of 1/10 volume of 11% formaldehyde in diluent (0.143 M NaCl, 1.43 mM EDTA, 71.43 mM HEPES-KOH) with gentle agitation for 2h. Cell pellets were washed twice with 10ml cold TBS (20mM Tris-HCl, pH 7.5, 150mM NaCl) and once with 10 ml cold FA lysis buffer (100mM Hepes-KOH, pH 7.5, 300 mM NaCl, 2 mM EDTA, 2% Triton X-100, 0.2% Na Deoxycholate)/0.1% SDS, frozen in liquid nitrogen and stored at −80°C. *S. pombe* cell pellets were resuspended in 400μL of cold 1x FA lysis buffer/0.5 % SDS containing 1x complete protease inhibitor cocktail (Roche) and 1mM PMSF and mixed with thawed *S. cerevisiae* pellet (approximately 100 ml cells OD600=0.4). ChIP and sequencing was performed as described above. Calculation of Occupancy Ratio (OR) and data analysis was performed as described in ^30^. The number of reads at each position were normalized to the total number of reads for each sample (RPM: Reads Per Million), multiplied by the occupancy ratio (OR) and shown in the Integrated Genome Viewer from the Broad Institute. Primers used for qPCR analysis are given in Table S3.

#### Immunofluorescence and microscopy

Indirect immunofluorescence to visualize spindles used a rat anti-tubulin antibody (AbD serotec) at a dilution of 1:50 and an anti-rat FITC conjugated antibody (Jackson Immunoresearch) at a dilution of 1:100. Cells were fixed in formaldehyde for visualization of TetR-GFP and Spc42-tdTomato foci. Yeast were mounted onto a glass slide mounted in Vectashield (Vector Laboratories, Peterborough UK) and imaged on a Zeiss Axio Imager Z1 equipped with a x100 α Plan Fluar/1.45 NA (oil) objective lens. Images were recorded using a Photometrics Evolve EMCCD camera (Photometrics, Tucson, USA) controlled using MicroManager 1.4 aquisition software (US National Institutes of Health). The fluorescent intensity and distance between the GFP foci were measured using a custom ImageJ plugin that can be found on the github repository https://github.com/dkelly604/CellClicker_.

#### RNA isolation and RNA-seq

Cell pellets were lysed by bead-beating in RLT buffer (Qiagen) and RNA was isolated using the RNeasy mini kit (Qiagen) according to the manufacturer’s instructions except that on-column DNA digestion was performed using the Qiagen DNase digestion kit. RNA concentration was determined by nanodrop. For cDNA synthesis for qRT-PCR, 12ng purified total DNA, diluted in HyClone dH2O and 10 mM Oligo(dT)15 primer (Roche) or 1.5 mM gene-specific reverse primer were incubated at 65°C for 10 min before placing on ice to denature RNA. Subsequently, 4 μl 5xTranscriptor RT reaction buffer (Roche), 0.5 μl RNase OUT (Fisher), 1 μM dNTPs and 0.5 μl Transcriptor reverse transcriptase plus Hyclone dH2O were added to 20 μl and incubated at 55 °C for 3 h before heat inactivation of Transcriptor Reverse Transcriptase at 85 °C for 5 min. RNA was depleted of rRNA and libraries prepared for sequencing by Genecore, EMBL. Sequencing was also performed by Genecore on an Illumina Next Seq 500 with a read l length of 75 and multiplexed with a pool size of 4.

#### Hi-C library preparation and data analysis

Hi-C protocol was modified from^31^and^32^. Cells were cultured, fixed and lysed as described in^31^. Briefly, 200 ml of cells at OD∼0.6 carrying *pMET-CDC20* were arrested in metaphase at 25 °C in the presence and absence of tension as described above, fixed with 3% formaldehyde for 20 minutes at 25 °C at 250 rpm and the reaction was quenched for 5 minutes by the addition of 0.35 M glycine (final concentration). Cells were washed with cold water, resuspended in 5 ml 1x NEBuffer 2 and frozen in liquid nitrogen. Lysates were prepared by grinding the frozen pellet in a chilled mortar with a pestle for 15 minutes and 1/10^th^ of the initial pellet weight (∼0.5 g) was taken for further processing. Restriction enzyme digestion (*Dpn*II), filling-in, ligation, crosslink reversal, DNA concentration and purification and biotin removal were carried out as described in ^32^. DNA was then fragmented on a Bioruptor Plus sonication device (Diagenode) for a total of 2x 30 cycles 30 seconds on/off at High setting. Following DNA end repair and A-tailing using T4 DNA polymerase, T4 Polynucleotide Kinase and Klenow fragment DNA polymerase I (as in ^32^), Hi-C libraries were fractionated using Ampure XP beads as previously described in ^31^. Biotin pull-down, adapter ligation (NextFlex, Bioo Scientific) and sequencing (EMBL Core Genomics Facility, Heidelberg, Germany) were carried out as in^32^. Hi-C read numbers are given in Table S4.

For Hi-C data analysis, Fastq reads were aligned to sacCer3 reference genome using HiC-Pro v2.11.1^33^ bowtie2 v2.3.4.1 (--very-sensitive -L 30 --score-min L,-0.6,-0.2 --end-to-end --reorder), removing singleton, multi hit and duplicated reads. Read pairs were assigned to restriction fragment (*Dpn*II) and invalid pairs filtered out. Valid interaction pairs were converted into the .cool contact matrix format using the cooler library, and matrixes balanced using Iterative correction down to one kilobase resolution. Multi-resolution cool files were uploaded onto a local HiGlass^34^ server for visualisation, cooler show was also used to generate individual plots for each chromosome. To generate pileups at centromeres/pericentromeric borders, the cooltools library was used, cool matrixes were binned at one kilobase resolutions. Plots were created around the midpoint of centromeres with ten/twenty-five/one-hundred kilobase flanks on each side, or around the midpoint of borders with forty kilobase flanks, showing the log10 mean interaction frequency using a colour map similar to HiGlass ‘fall’. All centromere/pericentromere annotations were duplicated in both the forward/reverse strand orientations to create a image which is mirror symmetrical. The observed over expected and ratio pile ups between samples were created in a similar fashion plotting the log2 difference between samples in the ‘coolwarm’ colour map, i.e. A/B; red signifying increased contacts in A relative to B and blue decreased contacts in B relative to A. Scripts are available at (https://github.com/danrobertson87/Paldi_2019).

## Data availability

ImageJ plugin to measure the fluorescent intensity and distance between the GFP foci can be found on the github repository https://github.com/dkelly604/CellClicker_. Scripts for Hi-C data analysis are available at https://github.com/danrobertson87/Paldi_2019.

## Acknowledgments

We are grateful to Bianka Baying and Vladimir Benes (Genecore, EMBL) for NGS and library preparation. We thank Robin Allshire, Stefan Bresson and David Tollervey for helpful discussions, Paul Megee for yeast strains and Weronika Borek, Stephen Hinshaw, Vasso Makrantoni and Pierre Romé for comments on the manuscript. This study was funded a Wellcome Senior Research Fellowship [107827] (AM, BA, FP), a Wellcome PhD studentship [109091] (FP), core funding for the Wellcome Centre for Cell biology [203149] (AM, FP, BA, DR, AK, DK), an ERC Consolidator Award [311336] (MJN, SS) and a Wellcome Trust Investigator Award [200843] (MJN, SS).

## Author contributions

AM conceived and supervised the study; FP and BA designed and performed experiments and analyzed the data with AM; DR, AK and SS performed bioinformatics; SS developed protocols; SS, MN and JB interpreted data. AM and FP wrote the paper, with input from all authors.

## Supplementary information

**Figure S1.**
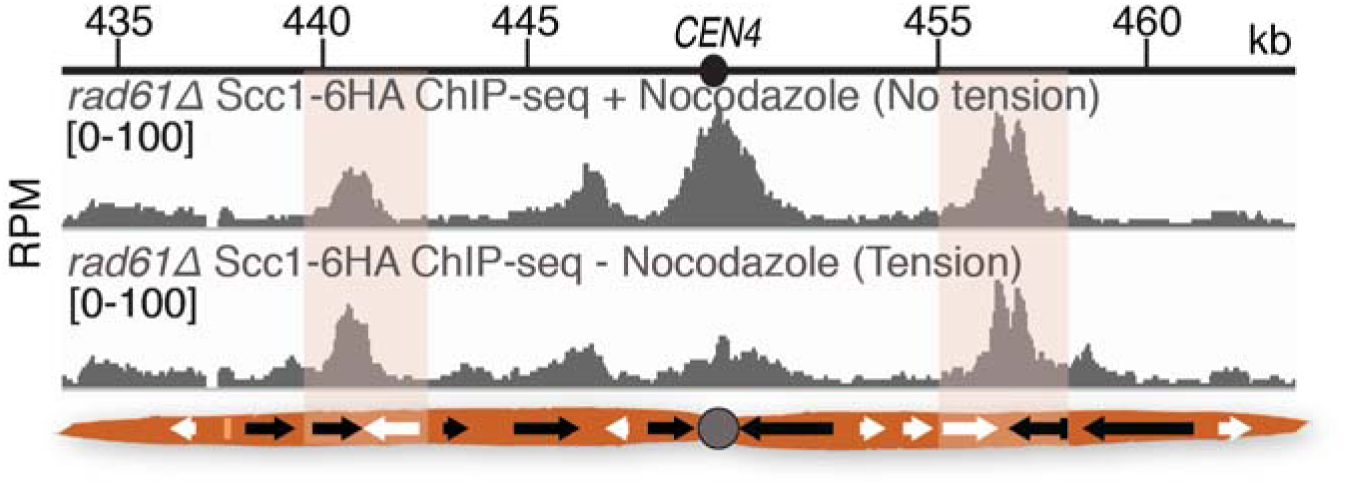
Wpl1/Rad61 is not required for the tension-dependent removal of cohesin at metaphase. Scc1-6HA calibrated ChIP-seq profiles for the pericentromeric region of chromosome IV are shown for *rad61Δ* cells arrested in metaphase, in the absence and presence of spindle tension.

**Figure S2.**
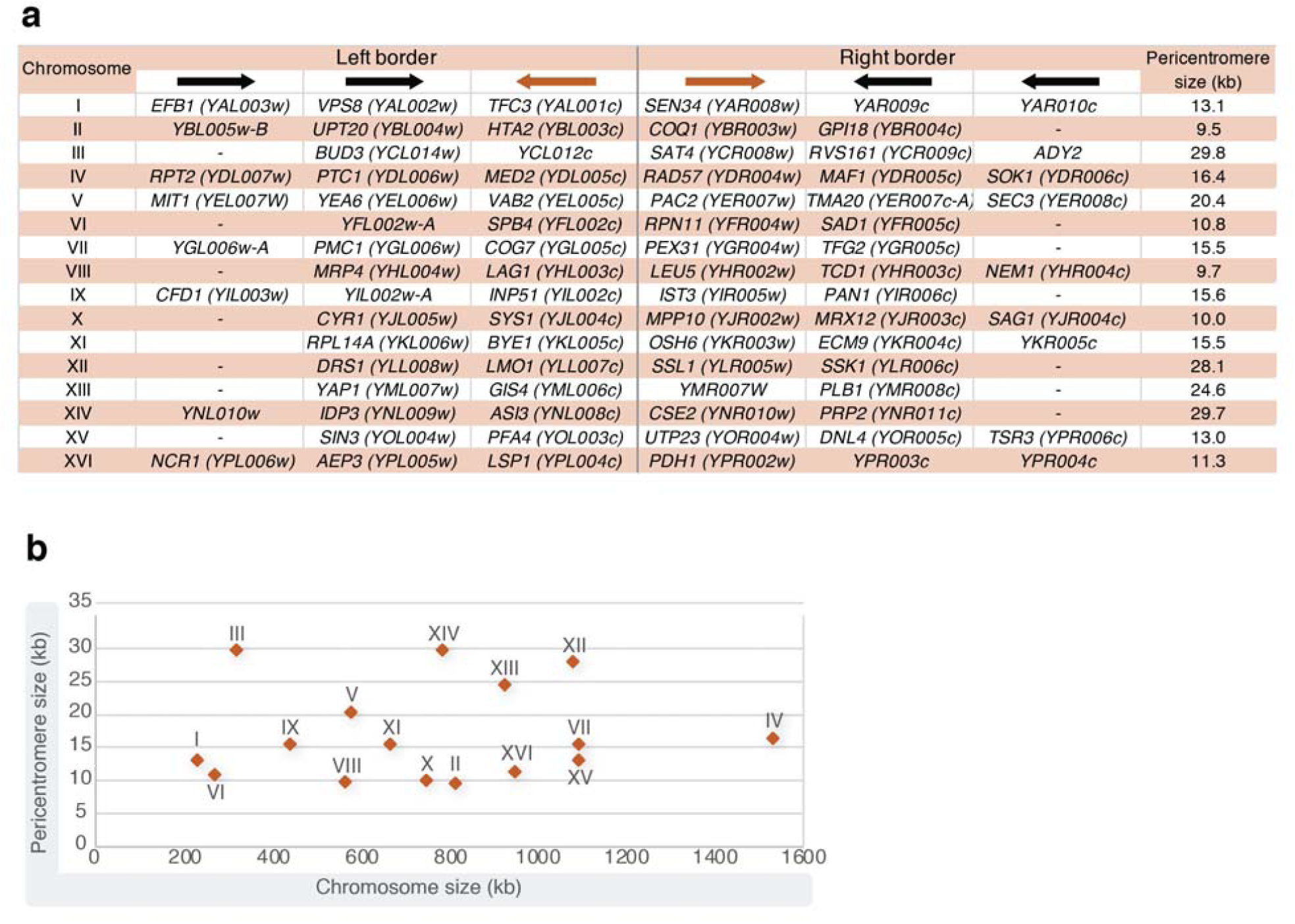
Genome overview of border gene organization and pericentromere size. **a**, Table of convergent genes identified at pericentromere borders for each chromosome, along with the corresponding pericentromere size. Borders were defined as the innermost cohesin peak near the centromere that persisted in the presence of tension. **b**, Pericentromere size determined in **a** plotted against chromosome size.

**Figure S3.**
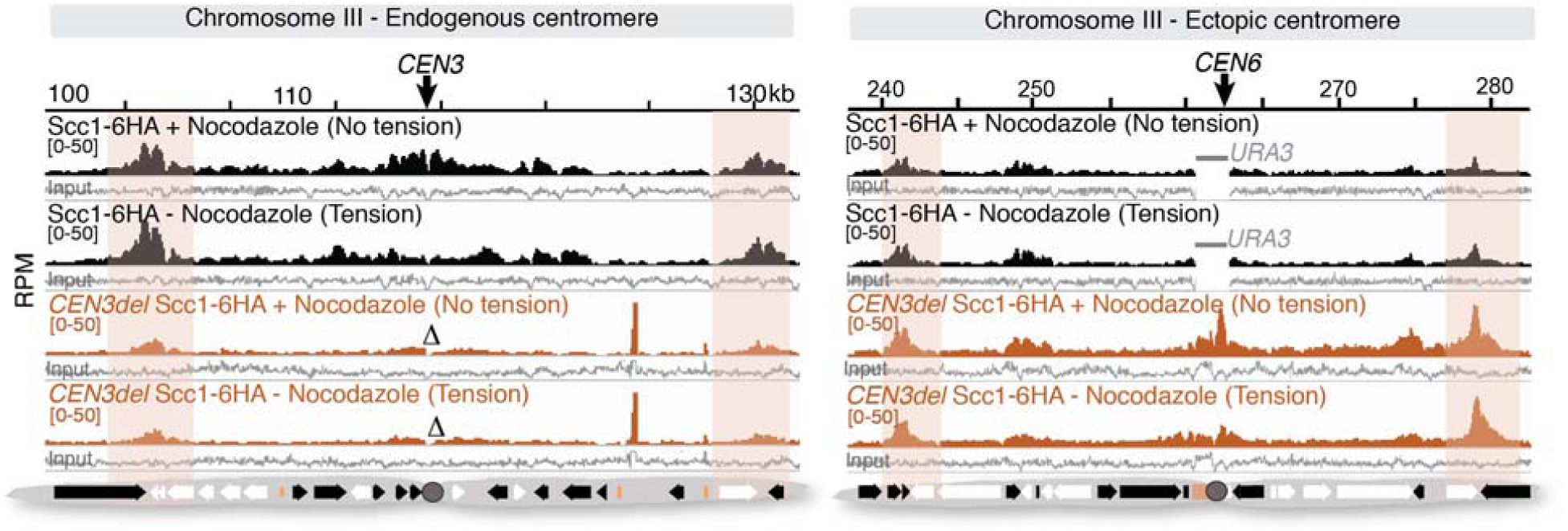
An ectopic centromere establishes new borders at convergent genes on a chromosome arm. Cohesin (Scc1) ChIP-Seq profiles for the region surrounding the endogenous centromere on chromosome III (left panel) and for a ∼50 Kb region of chromosome III surrounding the neo-centromeric arm site (right panel) are shown. Regions of tension-insensitive cohesin peaks at convergent sites flanking the endogenous and ectopic centromeres are highlighted.

**Figure S4.**
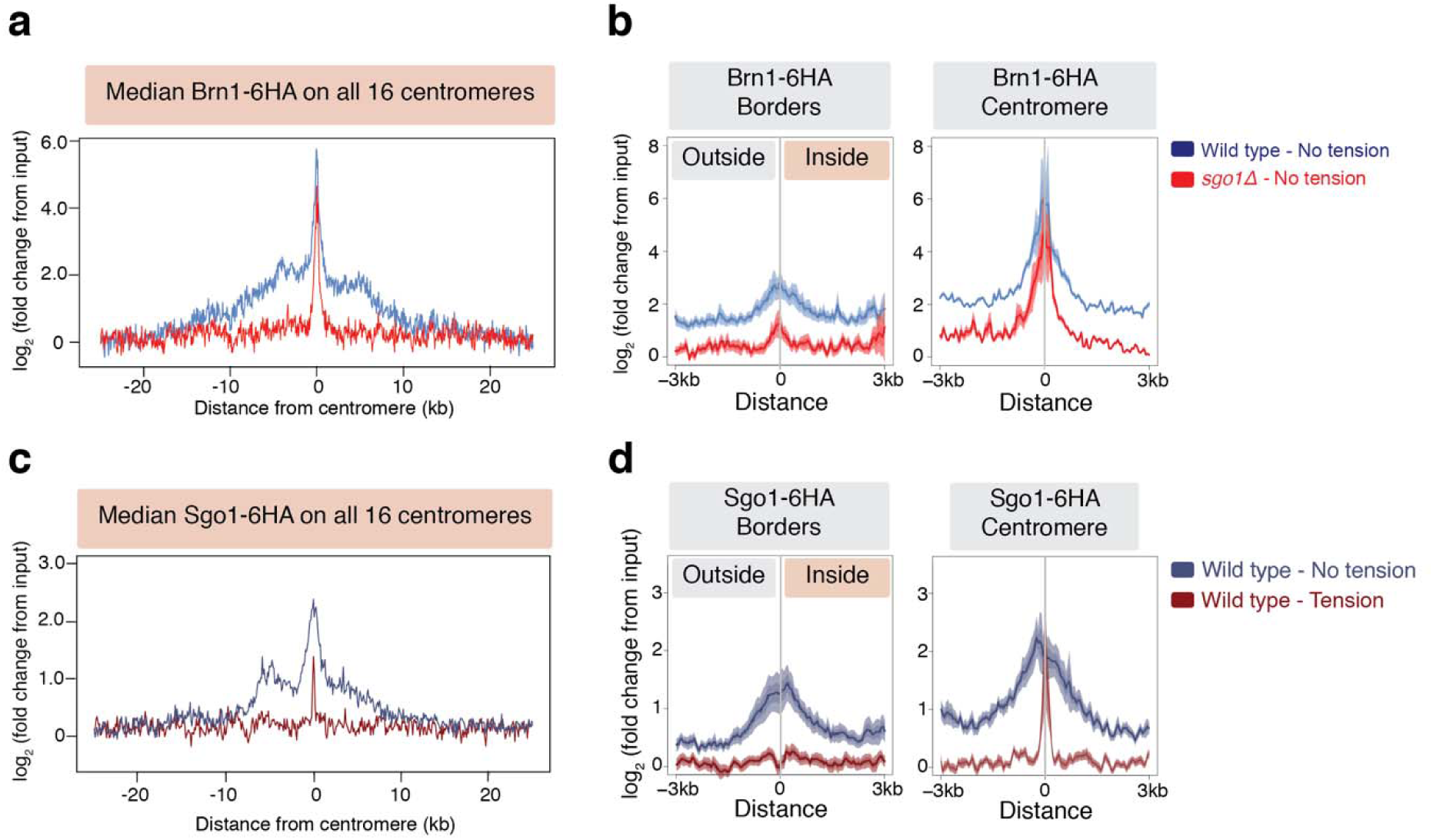
Shugoshin (Sgo1) enriches condensin at, and is removed in response to tension from, pericentromere borders. Condensin associates with pericentromere borders in a Sgo1-dependent manner in cells arrested in metaphase in the absence of tension. ChIP-Seq data used was previously published in^15^. **a**, Median condensin (Brn1) signal across a 50kb region surrounding all centromeres. **b**, Median Brn1 signal centered around borders (left panel) or centromere (right panel). Sgo1 is removed from the borders, but not core centromeres in response to spindle tension. **c**, Median Sgo1 enrichment by ChIP-Seq plotted over a 50kb region surrounding all centromeres in metaphase-arrested cells in the presence or absence of tension. **d**, Median Sgo1 signal centered around borders (left panel) or centromere (right panel).

**Figure S5.**
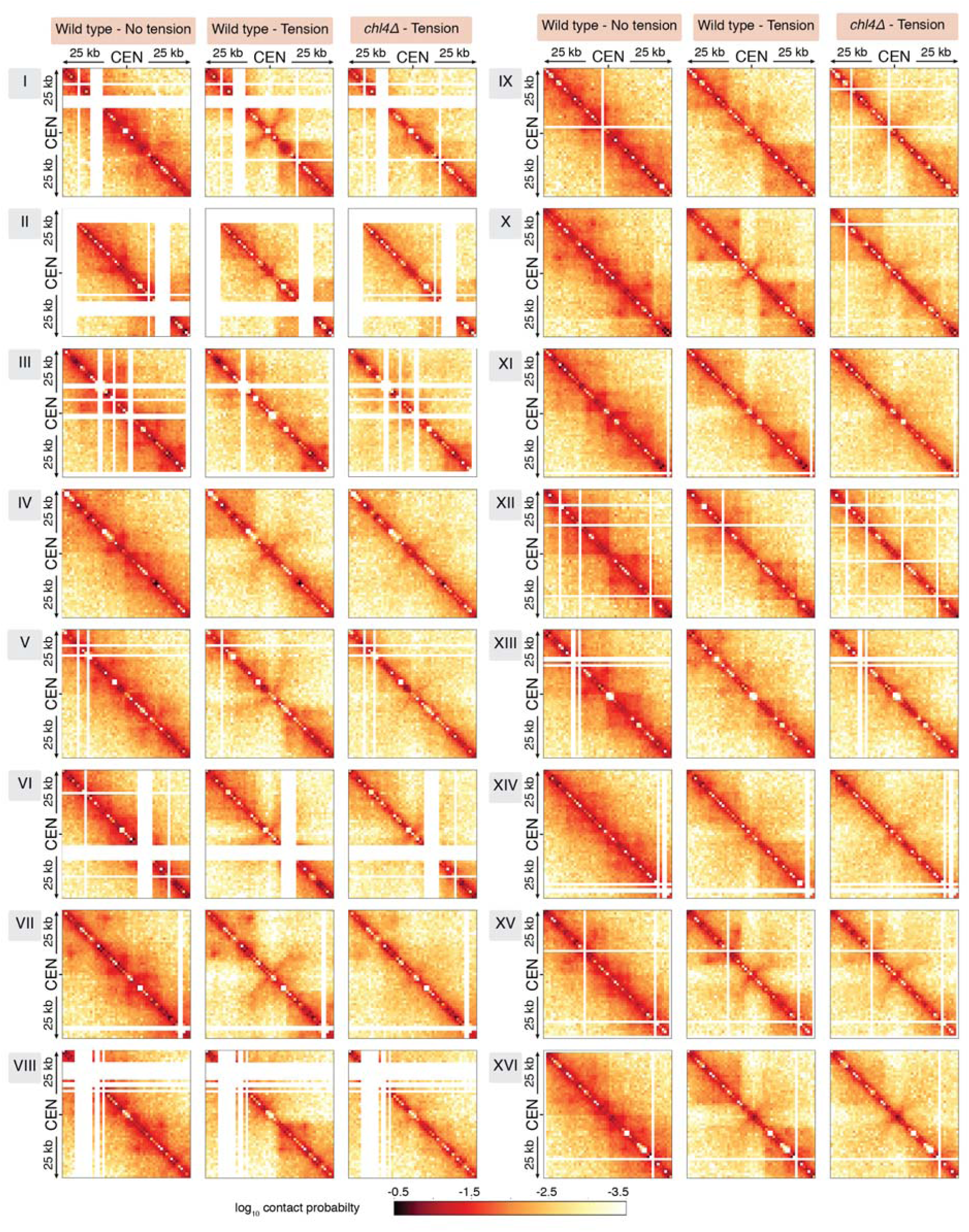
Changes in pericentromere structure on individual chromosomes in reponse to tension and in the absence of pericentromeric cohesin. Hi-C contact maps (1kb bin) over a 50kb region surrounding all centromeres in wild type cells without tension (left) and tension (middle), and in *chl4Δ* with tension (right).

**Figure S6.**
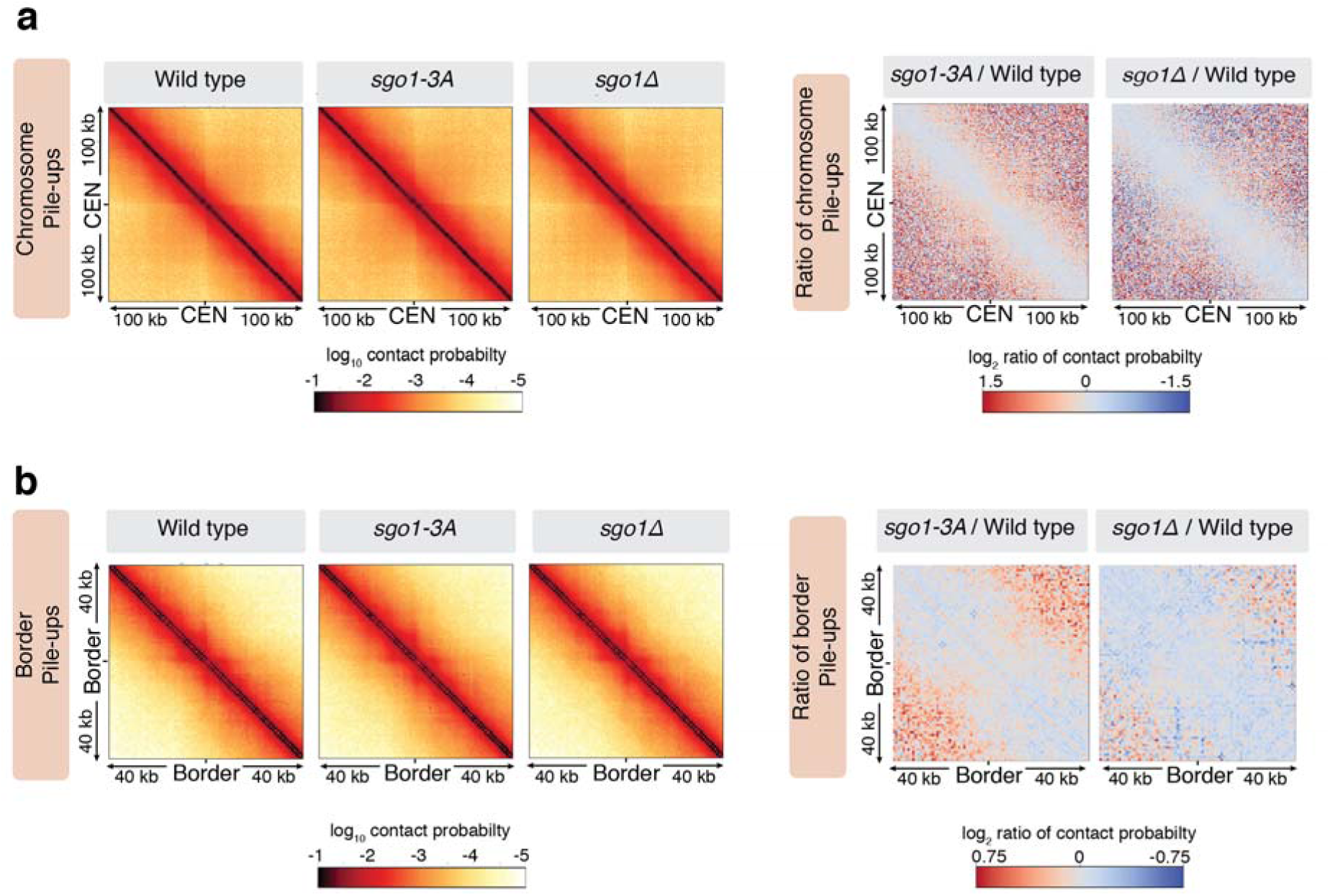
The absence of Sgo1 does not grossly alter pericentromere structure at metaphase without tension. Hi-C analysis of *sgo1-3A* and *sgo1Δ* in metaphase-arrested cells in the absence of tension reveals similar patterns to wild type. Data for wild type was reproduced from Figure 3a for comparison. **a**, Pile-ups (bin size 1kb) of *cis* contacts surrounding all 16 centromeres in absence of spindle tension for the indicated strains (left three panels) or Log2 difference maps between wild type and *sgo1-3A* or *sgo1Δ* (right two panels) detect little change. **b**, Pile-ups (bin size 1kb) and Log2 difference maps of *cis* contacts surrounding all 32 borders.

**Figure S7.**
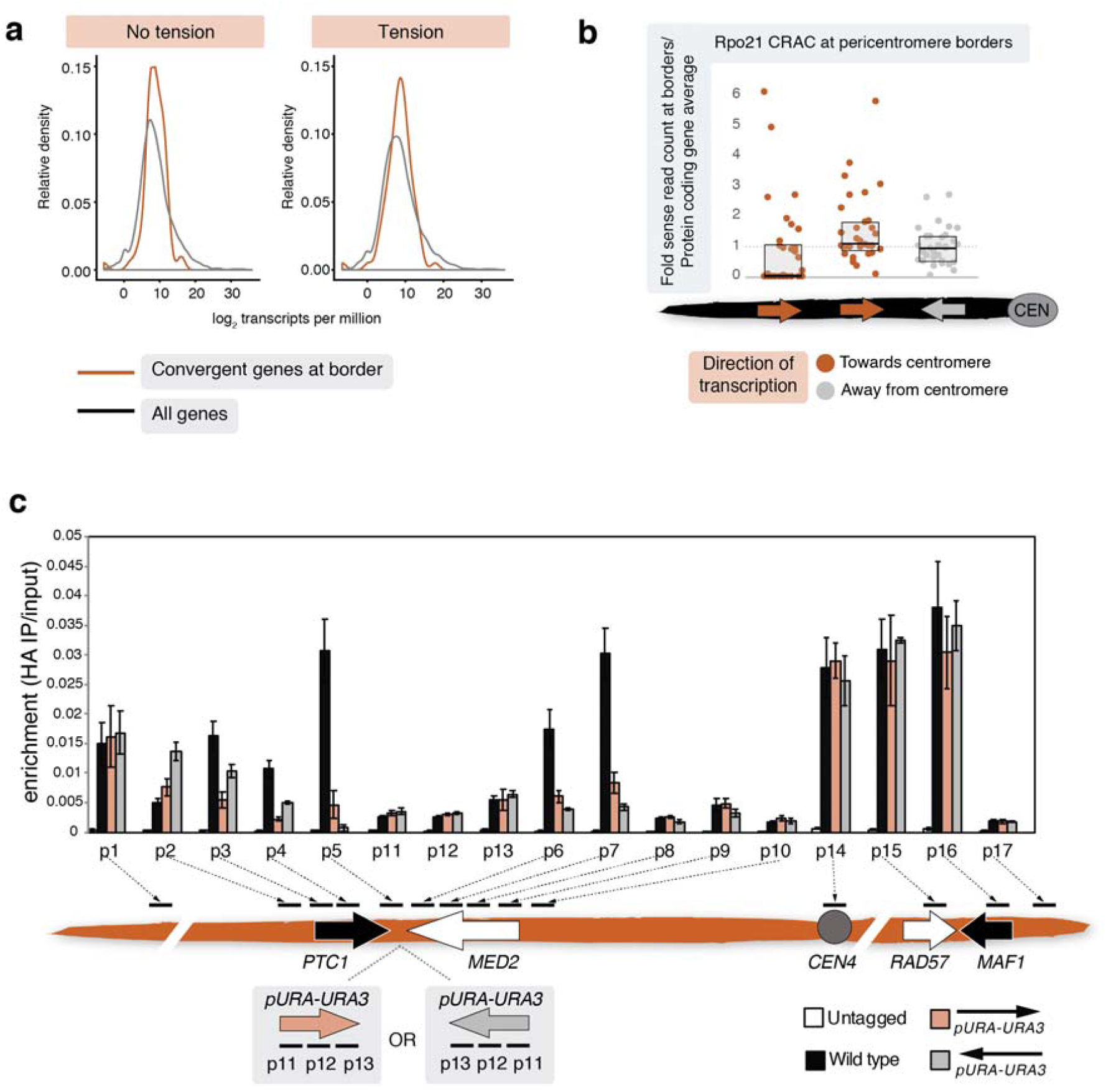
Transcription at pericentromere borders influences cohesin position. **a**, Genes at pericentromere borders are moderately transcribed on average. Relative RNA density for convergent gene pairs acting as borders compared to all genes is shown for no tension and tension conditions. Wild type cells were arrested in metaphase either in the presence or absence of nocodazole and RNA-Seq was performed. **b**, Boxplot of transcription levels of genes at pericentromere borders based on RNA polymerase II (Rpo21) Cross-linking and analysis of cDNA (CRAC) from^27^. Rpo21 CRAC sense read counts of genes at borders were normalized to the protein coding gene average and genes at pericentromere borders were grouped by their relative orientation to centromeres. Data points correspond to the mean of three separate repeats. **c**, Insertion of a *URA3* cassette between a convergent gene pair shifts the localization of cohesin in the direction of transcription. The *URA3* cassette was integrated in either orientation between the convergent gene pairs at the left pericentromere border on chromosome IV and cohesin (Scc1) ChIP-qPCR was performed. The positions of primers relative to the convergent genes and the *URA3* cassette are shown. Plot shows the mean of three biological replicates with error bars representing standard error.

**Figure S8.**
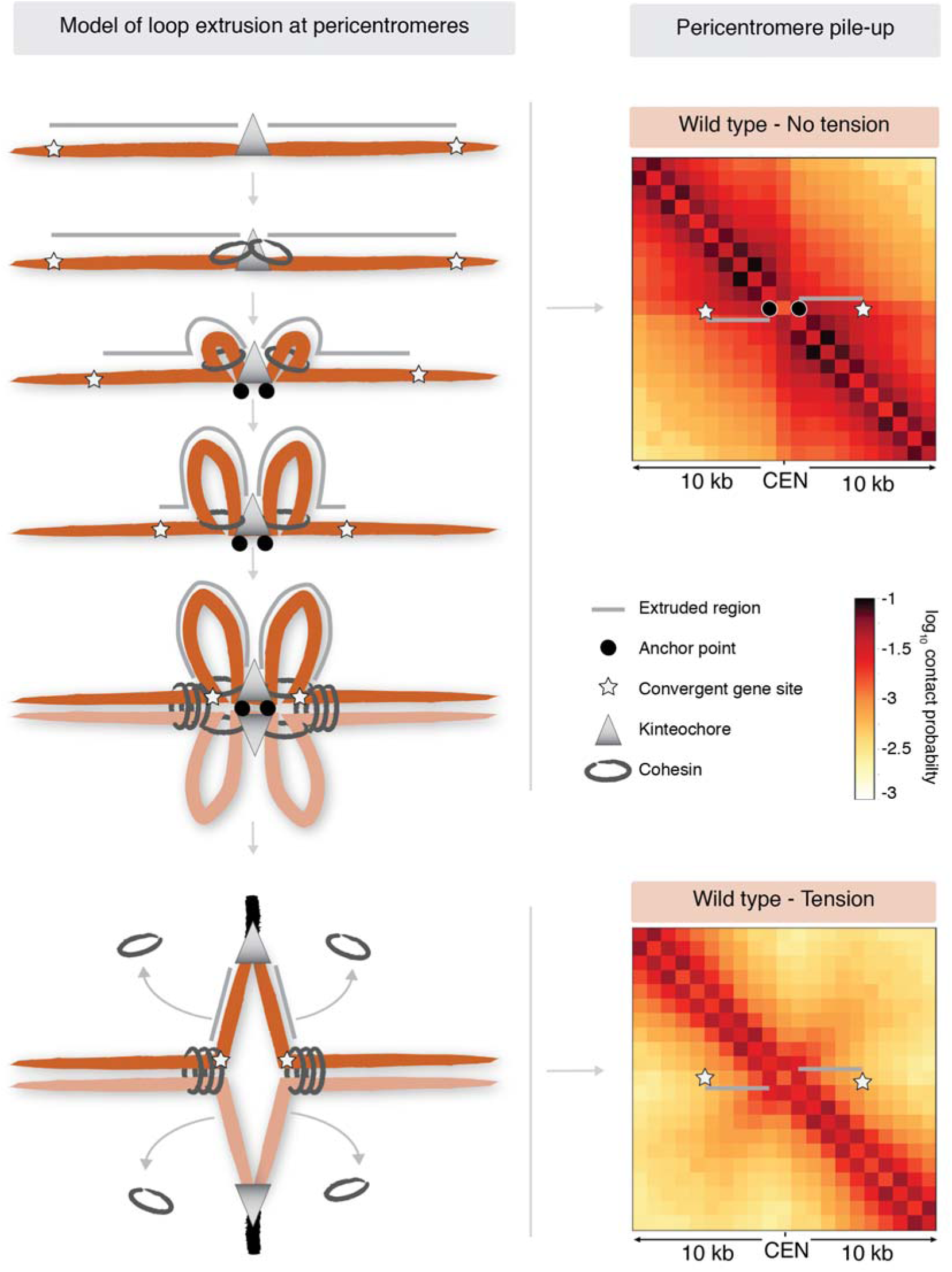
Model of loop extrusion at pericentromeres. Cohesin rings loaded at kinetochores make extrude a single chromatin loop at either side of the pericentromere until halted by a convergent gene site at pericentromere borders. Intramolecular cohesin at the base of loops is passively removed from chromosomes when biorientation extends pericentromeric chromatin outwards, converting centromere-flanking *cis*-loops to a V-shaped structure. Data for wild type was reproduced from Figure 3a, b.

**Table S1.** *Saccharomyces cerevisiae* strains used in this study.

| Strain | Relevant genotype | Figure |
| --- | --- | --- |
| AMy1105 | *MATa cdc20::URA3::pMET-CDC20 SCC1-6HA* | 1a, b, c, 1e, f, 2b, 3a, b, 3d, 4a, S3 |
| AMy1145 | *MATa SCC1-6HA* | S7C |
| AMy2508 | *MATa, cdc20::URA3::pMET-CDC20* | 3a, b, c, d, e, f, S5, S6a, b |
| AMy3950 | *MATa cdc20::URA3::pMET-CDC20 SCC1-6HA chl4Δ::KanMX6* | 1b, c, 3d |
| AMy6389 | *MATa cdc20::URA3::pMET-CDC20 SGO1-6HA::TRP1* | S4c, d |
| AMy6471 | *MATa, cdc20::URA3::pMET-CDC20 Spc42-tdTomato::NAT his3::PURA3::tetR-GFP::HIS3 ~1kbR_CEN3::tetOx224::URA3* | 2b, c |
| AMy6884 | *MATa cdc20::URA3::pMET-CDC20 SCC1-6HA sgo1(Y47A;Q50A;S52A)::hphMX4* | S6a, b |
| AMy7217 | *MATa SCC11-6HA cdc20::URA3::pMET-CDC20 SCC1-6HA sgo1Δ::KanMX6* | S6a, b |
| AMy14126 | *MATa, cdc20::URA3::pMET-CDC20 his3::PURA3::tetR-GFP::HIS3 SCC1-6HA::TRP1 SCC2-6HIS-3xFLAG::KAN ~1kbR_CEN3::tetOx224::URA3* | S7a |
| AMy16144 | *MATa cdc20::URA3::pMET-CDC20 CEN3Δ::LEU2 CEN6-URA3::CHRIII ~260kb SCC1-6HA* | S3 |
| AMy16541 | *MATa SCC1-6HA PTC1-pURA3-URA3-MED2* | S7c |
| AMy16721 | *MATa SCC1-6HA MED2-pURA3-URA3-PTC1* | S7c |
| AMy22078 | *MATa cdc20::URA3::pMET-CDC20 SCC1-6HA pMAF1-MAF1::loxp (reversed orientation) pPTC1-PTC1::loxp (reversed orientation) pRPT2-RPT2::loxp (reversed orientation) pSOK1-SOK1-loxp-KANMX-loxp (reversed orientation)* | 4a, b |
| AMy22900 | *MATa, cdc20::URA3::pMET-CDC20 Spc42-tdTomato::NAT leu2::tetR-GFP::LEU2 ~7kbR_CEN3::tetOx224::URA3* | 2b, c |
| AMy22936 | *MATa cdc20::URA3::pMET-CDC20 Spc42-tdTomato::NAT leu2::tetR-GFP::LEU2 ~3kbR_CEN3::tetOx224::URA3* | 2b, c |
| AMy23081 | *MATa, cdc20::URA3::pMET-CDC20 Spc42-tdTomato::NAT leu2::tetR-GFP::LEU2 ~3kbR_CEN1::tetOx224::URA3* | 2b, c |
| AMy23082 | *MATa, cdc20::URA3::pMET-CDC20 Spc42-tdTomato::NAT leu2::tetR-GFP::LEU2 ~1kbR_CEN1::tetOx224::URA3* | 2b, c |
| AMy23125 | *MATa, cdc20::URA3::pMET-CDC20 Spc42-tdTomato::NAT leu2::tetR-GFP::LEU2~7kbR_CEN1::tetOx224::URA3* | 2b, c |
| AMy23185 | *MATa, cdc20::URA3::pMET-CDC20 Spc42-tdTomato::NAT leu2::tetR-GFP::LEU2 ~18kbR_CEN3::tetOx224::URA3* | 2b, c |
| AMy25236 | *MATa, cdc20::URA3::pMET-CDC20 Spc42-tdTomato::NAT leu2::tetR-GFP::LEU2 ~23kbR_CEN3::tetOx224::URA3* | 2b, c |
| AMy25297 | *MATa, cdc20::URA3::pMET-CDC20 Spc42-tdTomato::NAT leu2::tetR-GFP::LEU2 ~8kbR_CEN1::tetOx224::URA3* | 2b, c |
| AMy25298 | *MATa cdc20::URA3::pMET-CDC20 SCC1-6HA pMAF1-MAF1::loxp (reversed orientation) pPTC1-PTC1::loxp (reversed orientation) pRPT2-RPT2::loxp (reversed orientation) pSOK1-SOK1-loxp-KANMX-loxp (reversed orientation) SGO1-6HIS-3FLAG::URA3* | 4b |
| AMy25299 | *MATa cdc20::URA3::pMET-CDC20 SCC1-6HA pMAF1-MAF1::loxp (reversed orientation) pPTC1-PTC1::loxp (reversed orientation) pRPT2-RPT2::loxp (reversed orientation) pSOK1-SOK1-loxp-KANMX-loxp (reversed orientation) BRN1-6HIS-3FLAG::NATMX6* | 4b |
| AMy25379 | *MATa cdc20::URA3::pMET-CDC20 SCC1-6HA BRN1-6HIS-3FLAG::NATMX6* | 1e, f |
| AMy25409 | *MATa cdc20::URA3::pMET-CDC20 SCC1-6HA SGO1-6HIS-3FLAG::URA3* | 1e, f |
| AMy25764 | *MATa, cdc20::URA3::pMET-CDC20 Spc42-tdTomato::NAT leu2::tetR-GFP::LEU2 ~21kbR_CEN3::tetOx224::URA3* | 2b, c |
| AMy26822 | *MATa, cdc20::URA3::pMET-CDC20 chl4Δ::KanMX6* | 3d, e, f, g, S5 |
| AMy26964 | *MATa, cdc20::URA3::pMET-CDC20 Spc42-tdTomato::NAT leu2::tetR-GFP::LEU2 ~12kbR_CEN1::tetOx224::URA3* | 2b, c |
| AMy26965 | *MATa, cdc20::URA3::pMET-CDC20 Spc42-tdTomato::NAT leu2::tetR-GFP::LEU2 ~12kbR_CEN3::tetOx224::URA3* | 2b, c |
| AMy26966 | *MATa, cdc20::URA3::pMET-CDC20 SCC1-6HA rad61Δ::TRP1* | S1 |
| AMy27213 | *MATa, cdc20::URA3::pMET-CDC20 Spc42-tdTomato::NAT leu2::tetR-GFP::LEU2 ~4kbR_CEN4::tetOx224::URA3* | 4c, d, e |
| AMy27214 | *MATa, cdc20::URA3::pMET-CDC20 Spc42-tdTomato::NAT leu2::tetR-GFP::LEU2 ~11.5kbR_CEN4::tetOx224::URA3* | 4c, d, e |
| AMy27215 | *MATa, cdc20::URA3::pMET-CDC20 Spc42-tdTomato::NAT leu2::tetR-GFP::LEU2 pMAF1-MAF1::loxp (reversed orientation) pPTC1-PTC1::loxp (reversed orientation) pRPT2-RPT2::loxp (reversed orientation) pSOK1-SOK1-loxp-KANMX-loxp (reversed orientation) ~13.5kbR_CEN4::tetOx224::URA3* | 4c, d, e |
| AMy27216 | *MATa, cdc20::URA3::pMET-CDC20 Spc42-tdTomato::NAT leu2::tetR-GFP::LEU2 pMAF1-MAF1::loxp (reversed orientation) pPTC1-PTC1::loxp (reversed orientation) pRPT2-RPT2::loxp (reversed orientation) pSOK1-SOK1-loxp-KANMX-loxp (reversed orientation) ~4kbR_CEN4::tetOx224::URA3* | 4c, d, e |

**Table S2.** Plasmids generated in this study.

| **Plasmid** | **Characteristics** |
| --- | --- |
| AMp1298 | *pMAF1-MAF2 LoxP-KanMX6-LoxP* |
| AMp1302 | *pPTC1-PTC1 LoxP-KanMX6-LoxP* |
| AMp1332 | *pRPT2-RPT2 LoxP-KanMX6-LoxP* |
| AMp1360 | *pSOK1-SOK1 LoxP-kanMX6-LoxP* |
| AMp1411 | *pRS306(tetOx224) + 502bp genomic sequence (2901bp right of centromere 3) to integrate tetOs ~3kb to right of CEN3* |
| AMp1412 | *pRS306(tetOx224) + 635bp genomic sequence (6453bp right of centromere 3) to integrate tetOs ~7kb to right of CEN3* |
| AMp1413 | *pRS306(tetOx224) + 514bp genomic sequence (17546bp right of centromere 3) to integrate tetOs ~18kb to right of CEN3* |
| AMp1433 | *pRS306(tetOx224) + 676bp genomic sequence (674bp right of centromere 1) to integrate tetOs ~1kb to right of CEN1* |
| AMp1436 | *pRS306(tetOx224) + 443bp genomic sequence (7123bp right of centromere 1) to integrate tetOs ~7kb to right of CEN1* |
| AMp1437 | *pRS306(tetOx224) + 377bp genomic sequence (3456bp right of centromere 1) to integrate tetOs ~3kb to right of CEN1* |
| AMp1538 | *pRS306(tetOx224) + 442bp genomic sequence (8105bp right of centromere 1) to integrate tetOs ~8kb to right of CEN1* |
| AMp1539 | *pRS306(tetOx224) + 632bp genomic sequence (23137bp right of centromere 3) to integrate tetOs ~18kb to right of CEN3* |
| AMp1562 | *pRS306(tetOx224) + 611bp genomic sequence (20810bp right of centromere 3) to integrate tetOs ~21kb to right of CEN3* |
| AMp1669 | *pRS306(tetOx224) + 505bp genomic sequence (12611bp right of centromere 1) to integrate tetOs ~12kb to right of CEN1* |
| AMp1670 | *pRS306(tetOx224) + 375bp genomic sequence (11156bp right of centromere 3) to integrate tetOs ~12kb to right of CEN3* |
| AMp1676 | *pRS306(tetOx224) + 535bp genomic sequence (3765bp right of centromere 4) to integrate tetOs ~4kb to right of CEN4* |
| AMp1677 | *pRS306(tetOx224) + 594bp genomic sequence (11426bp right of centromere 4) to integrate tetOs ~11kb to right of CEN4* |
| AMp1678 | *pRS306(tetOx224) + 560bp genomic sequence (13745bp right of centromere 4) to integrate tetOs ~14kb to right of CEN4 in MAF1-SOK1 reversed orientation strain* |

**Table S3.**
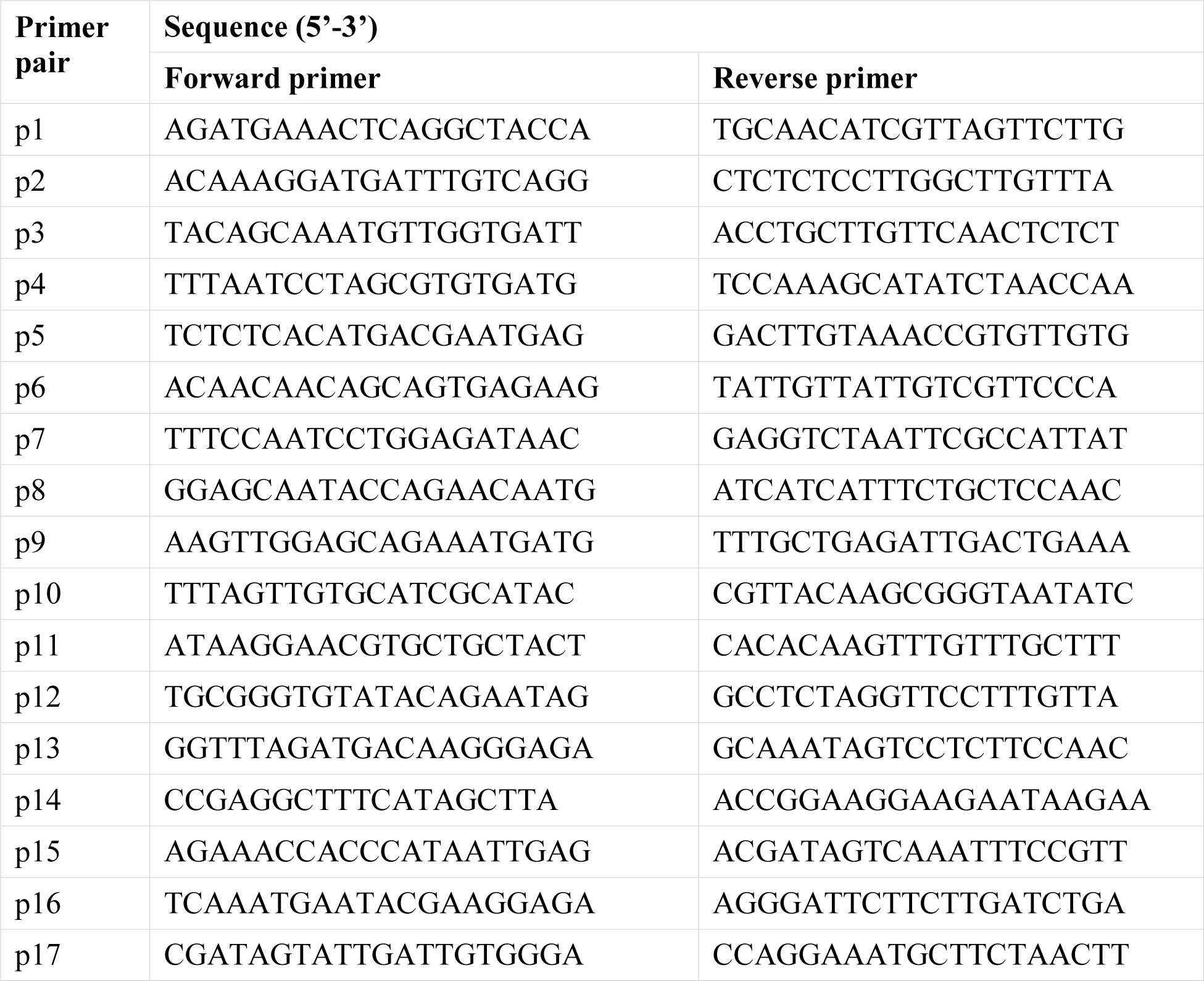
qPCR primer sequences used in this study.

**Table S4.**
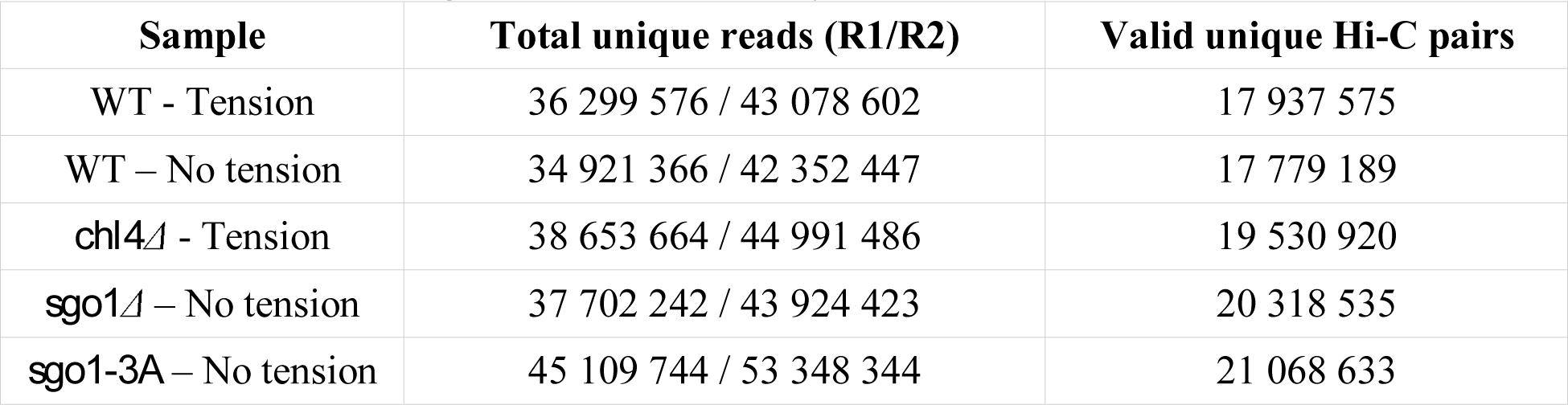
Hi-C libraries generated in this study.

| **Sample** | **Total unique reads (R1/R2)** | **Valid unique Hi-C pairs** |
| --- | --- | --- |
| WT - Tension | 36 299 576 / 43 078 602 | 17 937 575 |
| WT – No tension | 34 921 366 / 42 352 447 | 17 779 189 |
| *chl4Δ* - Tension | 38 653 664 / 44 991 486 | 19 530 920 |
| *sgo1Δ* – No tension | 37 702 242 / 43 924 423 | 20 318 535 |
| *sgo1-3A* – No tension | 45 109 744 / 53 348 344 | 21 068 633 |

